# Integrative Genomic and Functional Analyses Reveal NINL as a Modulator of Tau Aggregation

**DOI:** 10.64898/2025.12.12.694063

**Authors:** Samantha K. Swift, Guangming Huang, J. Nicholas Cochran, Miguel A. Minaya, Patricia A. Castruita, Katherine J. Miller, Emma Starr, Grant Galasso, Jacob A. Marsh, Aimee W. Kao, Jennifer S. Yokoyama, Celeste M. Karch

## Abstract

**Introduction:** Proteostasis dysfunction is a hallmark of frontotemporal dementia (FTD) and Alzheimer’s disease (AD), yet the genetic and molecular pathways that disrupt protein homeostasis remain poorly understood.

**Methods:** We integrated human genetics, transcriptomics, and functional studies to identify proteostasis network components involved in tauopathy.

**Results:** We identified 18 proteostasis network genes harboring 75 rare, damaging variants enriched in FTD and/or AD. These genes, spanning multiple proteostasis pathways, were differentially expressed in *MAPT* mutant neurons and dysregulated in FTD and AD brains. *NINL,* which encodes Nlp, emerged as the only gene consistently upregulated across all datasets. *NINL* overexpression reduced tau seeding and enhanced lysosomal proteolytic activity, whereas two FTD-enriched *NINL* frameshift variants impaired Nlp expression and abolished these protective effects.

**Discussion:** Our findings identify a set of proteostasis genes with genetic and transcriptional links to neurodegeneration and reveal *NINL* as a novel regulator of tau aggregation, potentially upregulated as an adaptive response to proteotoxic stress.

## 1. Introduction

Frontotemporal lobar degeneration (FTD) and Alzheimer’s disease (AD) are neurodegenerative disorders characterized by selective neuronal vulnerability and pathological tau aggregation. AD and a subset of FTD cases, known as frontotemporal lobar degeneration with tau inclusions (FTLD-tau), share the hallmark feature of tau aggregation in the brain and are collectively referred to as tauopathies. Although common genetic risk loci and causal mutations have been identified in tauopathies[1–7], a substantial proportion of disease heritability remains unexplained, suggesting that additional genetic factors, particularly rare variants, contribute to disease susceptibility. Defining these genetic contributors provides critical insight into the molecular mechanisms that drive neurodegeneration, revealing core pathogenic pathways and potential therapeutic targets.

Disruption of proteostasis is a feature of both FTD and AD. Impaired autophagy-lysosome function, defective chaperone activity, and the accumulation of aggregation-prone proteins, such as tau, converge to drive neuronal dysfunction and disease pathophysiology in FTD and AD [8–12]. While prior studies implicate proteostasis pathways in disease risk [13–20], the extent to which rare genetic variation affects components of the proteostasis network and how such variation translates into molecular dysfunction in human neurons and brain tissue remains poorly understood. Moreover, it is unclear whether specific proteostasis genes play direct, mechanistically actionable roles in modulating tau pathology.

By combining human genetics, disease model systems, and functional biology, our study aims to define proteostasis components that contribute to tau pathophysiology. Here, we integrated rare variant analyses from FTD and sporadic, early-onset AD cohorts with transcriptomic data from *MAPT* mutant iPSC-derived neurons and human tauopathy brain tissue to identify proteostasis network genes with coordinated genetic and transcriptional alterations. Across these datasets, we identify *NINL*, a motor-associated and trafficking-related protein, as consistently upregulated in tau-associated disease. Using functional assays, we validated *NINL* as a novel regulator of tau aggregation.

## 2. Materials and Methods

### 2.1 Whole genome sequencing and rare variant analysis

We performed secondary analyses of previously reported whole genome sequencing (WGS) data of early onset AD (EOAD), FTD, and control patients (**Supplemental Table 1**) [7]. Cases and controls were clinically evaluated as previously described[21, 22]. All participants or their surrogates provided written informed consent to participate in the study, and institutional review boards approved all aspects of the study.

Briefly, short-read, paired-end WGS was performed using Illumina HiSeq X. On average, 92% of bases were covered at a sequencing depth of 20x, along with a mean sequencing depth of 34x [7]. Sequencing reads were aligned to the reference genome hg19, and variants were called using GATK 3.8 [7]. Population frequency was determined using 1000 Genomes Phase 3, TOPMed Bravo, dbSNP (release 151), WGSA 0.7, which includes ExAC, gnomAD, ESP, and UK10K [23–28] Variants were filtered using SnpSift 4.3s, along with local and population frequency, predicted deleteriousness (annotated using CADD v1.3[29], and segmentation for function [7]. Variants analyzed had a maximum minor allele count of three (0.1% frequency in the cohort) and a minor allele frequency of 0.01% in population databases.

### 2.2 RNA sequencing analysis

RNA sequencing was performed on iPSC-derived neurons and human brains as previously described [30]. Briefly, samples were sequenced by an Illumina HiSeq 4000 Systems Technology with a read length of 1×150 bp and an average library size of 36.5 ± 12.2 million reads per sample. Salmon (v. 0.11.3) [31] was used to quantify gene expression (GRCh38.p13).

### 2.3 Cell Culture

HEK293T or HEK293T Tau Biosensor cells [32] were cultured in Dulbecco’s Modified Eagle Medium (DMEM; Gibco, Cat #: 61965-059), supplemented with 10% fetal bovine serum (FBS; Gibco, Cat #: 10270106) and 1% penicillin-streptomycin (Gibco, Cat #: 15140122). Cells were maintained in a humidified atmosphere with 5% CO_2_ at 37°C and were used within 5-15 passages. Cells were confirmed to be negative for mycoplasma contamination.

### 2.4 Plasmids and Transient Transfection

To evaluate the impact of *NINL* on cellular phenotypes, we used a pCMV6 plasmid containing *NINL* WT with a C-terminal myc-DDK-tag (OriGene, Cat #: RC206094). To model rare variants in *NINL*, we generated p.T275fs (c.823_824delAC) and p.R1202fs (c.3604dupA) by site directed mutagenesis in the pCMV6 *NINL* WT-Myc-DDK plasmid (Azenta). Mutagenized plasmids were confirmed by Sanger sequencing.

HEK293T or HEK293T-tau biosensor cells [32] were transiently transfected with pCMV6 *NINL* WT, p.T275fs, p.R1202fs, or an empty vector control using Lipofectamine 2000 following the manufacturer’s instructions (ThermoFisher, Cat #: 11668019). Briefly, cells were seeded in a Poly-D-Lysine (PDL; ThermoFisher, Cat #: A3890401) coated plate at 50% confluence. Plasmid (25ng) was mixed with 1ul of Lipofectamine 2000 in 1mL of serum-free Opti-MEM (Gibco, Cat #: 31985062) and incubated at room temperature for 15 minutes. Eighteen hours after adding the DNA/Lipofectamine complexes to the cells, the medium was replaced with fresh DMEM containing 10% FBS. Cells were utilized for downstream analysis at 24- or 48-hours post-transfection as indicated below.

### 2.5 Quantitative PCR (qPCR)

Gene expression was evaluated using qPCR via SYBR Green iTaq Universal SYBR Green Supermix (Bio-Rad, Cat #: 1725121). Primers specific to *NINL* included 5’-ACCTGGGATTCTGAGGACTTTG-3’ and 5’-ACTTTGCCGTCTCCGTCTTGAT-3’. Technical replicates were performed for each sample. *NINL* assays were run in separate wells from the housekeeping gene, *GAPDH*, to prevent interference. Primers for *GAPDH* included 5’-TGCACCACCAACTGCTTAGC-3’ and 5’-GGCATGGACTGTGGTCATGAG-3’. Data was analyzed via the comparative C_T_ method. Relative expression was quantified using the comparative Ct method. *NINL* Ct values were normalized to *GAPDH* for each sample and then compared to the mean normalized expression of the control cohort. Only samples for which the standard error was less than 20% between technical replicates were analyzed.

### 2.6 LysoTracker and LysoSensor

Acidic vesicles were measured by flow cytometry using LysoTracker and LysoSensor. HEK293T cells were transiently transfected with control or *NINL* plasmids for 48 hours using Lipofectamine 2000 (ThermoFisher, Cat #: 11668019) as described above. Cells were then incubated with LysoTracker Red (ThermoFisher, Cat #: L7528) at a dilution of 1:10,000 in DMEM supplemented with 10% FBS for 25 minutes at 37°C. LysoSensor Green (1:1,000; ThermoFisher, Cat #: L7535) was added to the cells for 60 seconds at the end of the LysoTracker incubation. Cells were then rinsed with PBS and collected via trypsinization (0.05%). The live cell population was labeled with Far Red Live/ Dead (ThermoFisher, Cat #: L34974) stain at 1:1,000 in PBS for 25 minutes. Flow cytometry was performed using a BD FACSymphony™ A1. LysoTracker Red was measured using a 405 nm laser with a BV480 filter, LysoSensor Green was measured using a 561 nm laser and a PE filter, and Far-Red Live/Dead was measured using a 637 nm laser and an APC filter. We analyzed 10,000 cells per replicate. Data were processed using FlowJo software (v10).

### 2.7 Immunoblotting

Immunoblotting was performed to measure expression of proteins in the autophagy-lysosome pathway. HEK293T cells were transiently transfected with control or *NINL* plasmids for 48 hours using Lipofectamine 2000 (ThermoFisher, Cat #: 11668019) as described above. Cells were harvested using trypsinization (0.05%), washed with ice-cold 1x PBS containing a protease inhibitor cocktail (ThermoFisher, Cat #: P2714), and centrifuged at 500 x g for 5 minutes at 4°C. Cells were lysed by sonication in RIPA buffer containing 50 mM Tris (pH 7.4), 150 mM NaCl, 1% Triton X-100, 1% sodium deoxycholate, 0.1% SDS, 50 mM NaF, 1 mM Na2VO4, 5mM EDTA, 1x phosphatase inhibitor (Sigma-Aldrich, Cat #: 4906845001), and protease inhibitor cocktail (1:500; Sigma-Aldrich, Cat #: P2714). Lysates were then centrifuged at 16000*x*g for 10 minutes at 4°C and transferred to a fresh tube. Protein concentration was determined using BCA (ThermoFisher, Cat #: 23225). Equal amounts of total protein (15 ug) were incubated at 70°C for 10 minutes in SDS sample buffer (ThermoFisher, Cat #: B0007) containing 10% β-mercaptoethanol. Proteins were resolved by SDS-PAGE (ThermoFisher, Cat #: NW04120BOX) and transferred onto a 0.2μm PVDF membrane (Millipore Sigma, Cat #: ISEQ00010). Membranes were blocked with 3% BSA in PBS-T (phosphate buffered saline with 0.1% TWEEN20) and incubated with primary antibodies overnight at 4°C. Primary antibodies used in this study include Nlp (Sigma-Aldrich, Cat #: HPA000686), Lamp1 (Cell Signaling, Cat #: 9091), LC3B (Cell Signaling, Cat #: 2775), 9E10 C-Myc (Sigma-Aldrich, Cat #: M4439), and β-actin (Cell Signaling, Cat #: 4970). After rinsing in PBS-T, membranes were incubated with horseradish peroxidase-conjugated secondary antibodies for one hour at room temperature. Secondary antibodies used in this study include anti-mouse IgG, HRP-linked Antibody (Cell Signaling, Cat #: 7076S) or anti-rabbit IgG, HRP-linked Antibody (Cell Signaling, Cat #: 7074S). Blots were visualized using Lumigen ECL Ultra (TMa-6) chemiluminescent substrate (Fisher Scientific, Cat #: NC0240697) on a Bio-Rad ChemiDoc Imaging System.

### 2.8 Measuring Autophagic Vesicles

Autophagic vesicle size and number were measured by confocal microscopy using CytoID Autophagy detection kit (Enzo, Cat #: ENZ-51031)[33]. Twenty-four hours after transfection, cells were replated at a density of 200,000 cells per well into 35mm live imaging dishes with a 20mm glass bottom well (Cellvis, Cat #: D35-20-1.5-N) coated with Poly-D-Lysine. Cells were grown in imaging dishes for an additional 24 hours. Two hours prior to imaging, cells were treated with ViaFluor Live Cell Microtubule Stain (Biotium, Cat #: 70063), and Verapamil HCl (Enzo, Cat #: ENZ-51031) was added at a final concentration of 10μm according to the manufacturer’s instructions. CytoID Green Reagent (Enzo, Cat #: ENZ-51031) and Hoechst 33342 Nuclear Stain (Enzo, Cat #: ENZ-51031) was added 30 minutes prior to imaging per the manufacturer’s instructions. In a subset of wells, cells were treated with 500nM Rapamycin and 10μM Chloroquine for 6 hours prior to imaging to stimulate the formation of autophagosomes and impair autolysosome fusion. Wells were imaged on a Nikon AX-R with NSPARC confocal microscope using a 60x oil immersion objective. Approximately 15 cells were imaged per well per condition. Images were analyzed using Imaris.

### 2.9 Monitoring Protease Activity

Substrate uptake and degradation by proteases were measured via live imaging using DQ-BSA [34, 35]. Twenty-four hours after transient transfection with control or *NINL* plasmids, HEK293T cells were replated at a density of 10,000 cells per well into a 96 well plate coated with Poly-D-Lysine. Cells were treated with DQ-BSA Red (1:1,000; ThermoFisher, Cat #: D12051) 24 hours after replating and imaged every 2 hours for 48 hours in a Sartorius Incucyte S3. Five images per well were acquired using a 10x objective. Images were analyzed using Incucyte analysis software.

To measure Cathepsin B activity, HEK293T cells were transiently transfected with control or *NINL* plasmids and replated at a density of 10,000 cells per well into a 96 well plate 24 hours later. Cells were treated with 1x Magic Red following the manufacturer’s instructions (Antibodies Inc., Cat #: 938) 24 hours after replating. Cells were imaged hourly for 12 hours in a Sartorius IncuCyte S3. Five images per well were taken using a 10x objective. Images were analyzed using IncuCyte analysis software.

### 2.10 Tau seeding activity

To investigate whether changes in *NINL* expression modulate tau seeding activity, we utilized a HEK293T tau biosensor cell line stably expressing tau repeat domain (RD) with the FTD-linked P301S variant of tau fused to either CFP or YFP [32, 36]. Cells were cultured in a 12 well plate. Twenty-four hours after transient transfection with control or *NINL* plasmids, cells were treated with 10 µM of human recombinant Tau-441 (2N4R) wild-type protein pre-formed fibrils (StressMarq, Cat #: SPR-471E) incubated with Lipofectamine 2000. After an additional 24 hours, cells were harvested using 0.05% trypsin, fixed in 4% paraformaldehyde for 10 minutes, and resuspended in flow cytometry buffer (2% FBS, 0.5% BSA in PBS). Flow cytometry was conducted using a BD LSR Fortessa™ Cell Analyzer (BD Biosciences). CFP and FRET were measured using a 405 nm laser with emissions collected at 405/50 nm and 525/50 nm, respectively. YFP fluorescence was measured using a 488 nm laser with emission at 525/50 nm. FRET signals were quantified as previously described [36], using cells expressing only RD-CFP or RD-YFP for spectral compensation. A bivariate plot of FRET vs. CFP was generated, and FRET-positive cells were identified using a triangular gating strategy. This gate was calibrated using biosensor cells treated with Lipofectamine alone as FRET-negative controls. For each experimental condition, 20,000 cells per replicate were analyzed. Data were processed using FlowJo software (v10).

## 3. Results

### 3.1 Rare variants occurring in proteostasis genes are enriched in FTD and AD

FTD and AD exhibit a high degree of heritability that remains unexplained, suggesting additional genetic factors likely contribute to the disease that have not yet been identified [3, 37]. Defining how rare variants modify FTD and AD risk can reveal core pathogenic pathways and therapeutic targets, while reciprocal analyses (e.g. testing for enrichment of rare variants within these pathways) enable a deeper understanding of genetic convergence in disease. Given the genetic and molecular evidence for the role of proteostasis genes in FTD and AD risk [10, 11, 14, 16], we sought to determine whether rare variants enriched in FTD and AD occur in genes involved in the proteostasis network. Here, we leveraged summary statistics from our prior study of rare variant enrichment performed in unrelated FTD cases (n=208), early onset AD cases (n=227), and healthy adult controls (n=671) [7] (**Supplemental Table 1**). We specifically asked whether proteostasis network genes, defined using the Proteostasis Consortium [38, 39] and prior literature [40], harbored rare variants enriched in FTD/AD cases. Rare variants were defined as those with a minor allele frequency of 0.0001 in all populations. Priority was given to variants present in 3 or more cases; present in 0 or 1 controls; and with a CADD score > 15 (**Figure 1A**). CADD scores above 15 are predicted to be damaging to the protein [29]. We identified 75 variants in 18 proteostasis network genes with rare variants that were enriched in FTD and/or AD cases and predicted to be damaging (**Figure 1B**). Across the 18 proteostasis genes, the 75 unique variants were identified and include missense, frameshift, splicing, and stop gain variants (**Supplemental Table 2**). The 18 proteostasis genes are involved in regulation of chaperone functions, autophagy-lysosome pathway, autophagosome and lysosome positioning, ubiquitin proteosome system, and stress response, among others (**Figure 1C**). Together, we show that rare variants involved in proteostasis network genes occur in FTD and AD.

**Figure 1.**
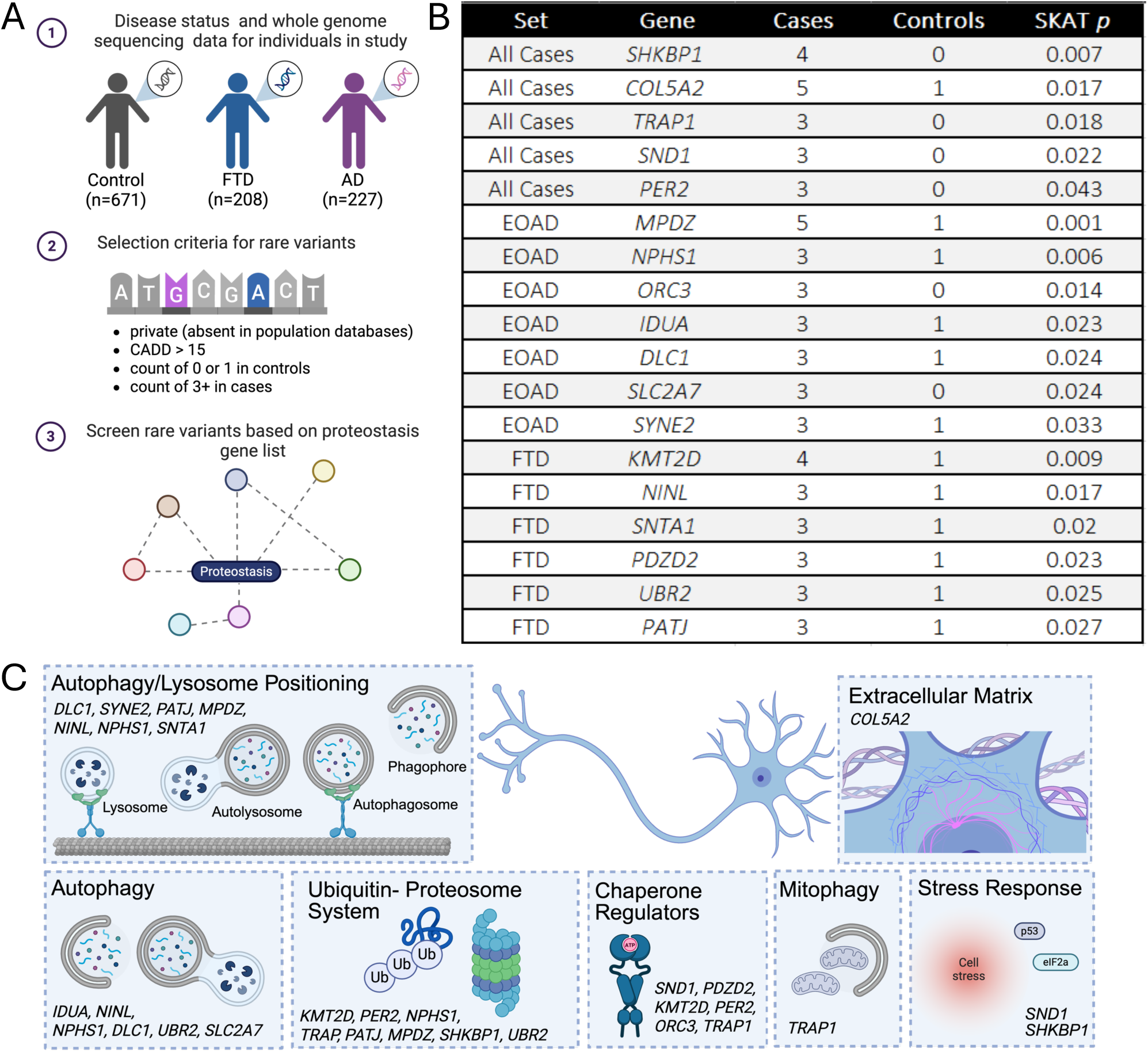
Rare variants in proteostasis associated genes are identified in AD and FTD cases. A. Schematic depicting workflow of variant identification from whole genome sequencing from Cochran et al [7]. FTD, Frontotemporal Dementia. AD, Alzheimer’s Disease. B. Table of genes that fall within proteostasis related pathways and that contain a rare variant that are enriched in FTD and AD. C. Diagram of pathways involving proteostasis related genes.

### 3.2 Proteostasis gene expression is altered in tauopathy

We next sought to determine whether proteostasis network genes harboring rare variants associated with FTD and AD also exhibit altered expression in disease models. This approach provides functional context for genetic risk, which could reveal how sequence variation translates into dysregulated molecular pathways. A subset of FTD and AD share a common pathological feature, tau aggregation. Thus, we analyzed transcriptomic data from human iPSC-derived neurons harboring *MAPT* mutations, which have been established as a powerful cellular system to model early stage tauopathy [30, 41–43]. Three *MAPT* mutations were analyzed along with their isogenic controls: p.P301L (exon 10 point mutation), IVS10+16 (splicing), and p.R406W (C-terminal point mutation) (**Figure 2A**)[30]. One gene, *SLC2A7*, was not expressed in the iPSC-derived neurons. Of the remaining 17 genes, 14 were differentially expressed in *MAPT* p.P301L neurons, and 10 were differentially expressed in *MAPT* IVS10+16 neurons, which is more than we would expect by chance (p= 0.0127 and 0.0003, respectively; **Figure 2B**). Only two of the 17 genes were differentially expressed in *MAPT* p.R406W neurons: *NINL* and *IDUA* (**Figure 2B**). Nine of the 18 genes were significantly differentially expressed in at least 2 of the 3 *MAPT* mutations (p<0.05; **Figure 2B**; **Supplemental Table 3**). Strikingly, one gene, *NINL,* exhibited increased expression in all three *MAPT* mutations (**Figure 2B**). These results demonstrate that proteostasis genes containing rare variants enriched in FTD and AD are altered in a stem cell model of tauopathy, suggesting these genes are more broadly related to pathways associated with tau pathophysiology.

**Figure 2.**
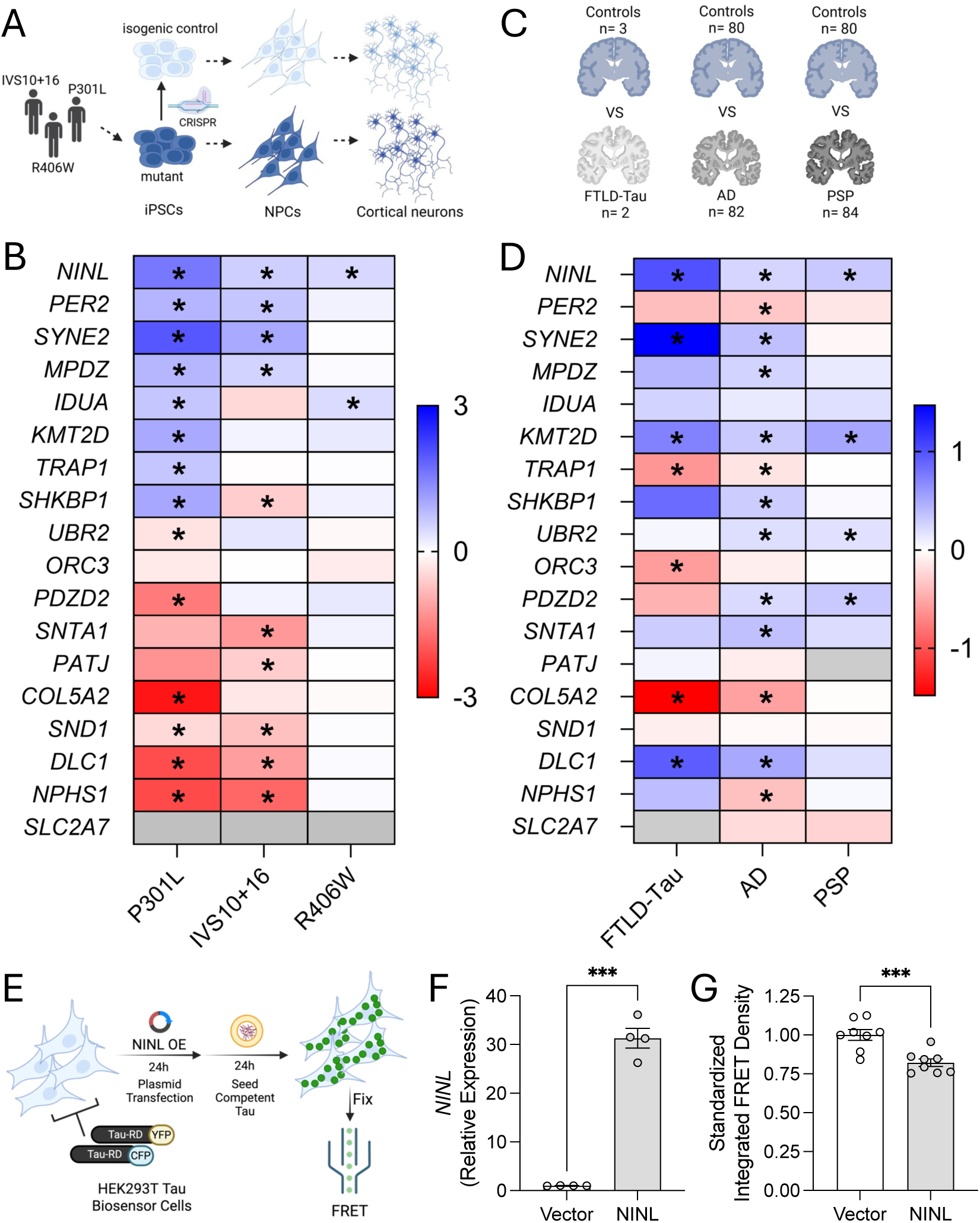
Proteostasis genes are differentially expressed in models of tauopathy. A. Schematic. FTD-associated *MAPT* mutant neurons and isogenic controls were differentiated from iPSC and subjected to RNAseq. B. Heatmap of logfold change. Differential gene expression data from *MAPT* mutant neurons. The 18 genes identified via whole genome sequencing (Figure 1B) are shown. C. Schematic depicting tauopathy (FTLD-Tau, AD, and PSP) and control brains. D. Heatmap. Differential gene expression data from human brains. The 18 genes identified via whole genome sequencing (Figure 1B) are shown. E-G. FRET-based tau seeding assay in HEK293T tau biosensor cells. E. Schematic. F. qPCR for *NINL* RNA. ***, p= 0.0006. Student’s t-test. G. Integrated FRET Density (FRET MFI x percent FRET positive cells) normalized to vector controls, ***, p=0.0009. Student’s t-test. Graphs represent mean ± SEM. Results represent 2 independent experiments with 4 replicates per condition in each experiment.

Due to their immaturity, human iPSC neurons likely capture early events associated with disease pathophysiology prior to protein aggregation [30, 41, 44–46]. To determine whether the 18 proteostasis network genes were associated with pathological events late in disease and in the presence of tau aggregates, we used transcriptomic data from FTLD-tau patients carrying a *MAPT* IVS10+16 mutation (n=2) and controls (n=3; **Figure 2C**). In the brains of FTLD-tau patients, *SLC2A7* was not detected, and 7 of the 17 remaining genes were differentially expressed (**Figure 2D**; **Supplemental Table 4**). To determine whether the 18 proteostasis network genes were associated with AD, we analyzed transcriptomic data from AD (n=82) and control (n=84) brains (**Figure 2C**). We identified 13 differentially expressed genes in AD brains (**Figure 2D**; **Supplemental Table 4**). *NINL*, *SYNE2*, *KMT2D*, *TRAP1*, *COL5A2*, and *DLC1* exhibit consistently significantly differential expression across FTLD-tau and AD brains (**Figure 2D**; **Supplemental Table 4**). To determine whether the 18 proteostasis network genes were altered in other forms of tauopathy, we examined transcriptomic data in brains from the primary, sporadic tauopathy, PSP (n=84; **Figure 2C**). In PSP brains, we identified 4 differentially expressed proteostasis network genes (of the 17 genes present in the dataset; **Figure 2D**; **Supplemental Table 4**). *UBR2* and *PDZD2* were significantly upregulated in PSP and AD brains (**Figure 2D**). *NINL* and *KMT2D* were significantly upregulated in FTLD-tau, AD, and PSP brains (**Figure 2D**). Together, we show that proteostasis network genes with genetic associations to FTD and AD are more broadly dysregulated at disease end stage.

### 3.3 NINL reduces tau seeding

We identified a single gene that contains rare variants enriched in FTD and that is significantly elevated across *MAPT* mutant neurons as well as in FTLD-Tau, PSP, and AD brains: *NINL*. Given this observation, we asked whether *NINL* plays a role in a central feature of tau pathophysiology: tau aggregation. To determine whether *NINL* expression impacts the propensity of tau to seed new aggregates, we used an engineered HEK293T line that stably expresses two constructs encoding a mutagenized form of the tau repeat domain (RD-P301S) conjugated to either CFP or YFP[32] (i.e. tau biosensor cells). Adding tau aggregates to these cultures has been shown to cause nucleation of the endogenous tau reporter proteins and aggregation that can be measured by FRET using flow cytometry [36] (**Figure 2E**). Surprisingly, overexpression of *NINL* wild-type (WT) in tau biosensor cells (**Figure 2F**) resulted in a significant reduction in tau seeding compared with vector controls (**Figure 2G**). Thus, *NINL* dampens tau seeding capacity, reducing tau aggregation. Considering these findings along with the observed increase in *NINL* in *MAPT* mutant neurons and tauopathy brains, we hypothesize that *NINL* is upregulated as an adaptive protective response to pathological stress.

### 3.4 *NINL* impacts lysosomal load

*NINL* encodes Nlp which is a motor-associated protein commonly known for its role in the centrosome and cell division [47–49]. In neurons, Nlp supports vesicle trafficking by interacting with motor complexes, such as dynein and dynactin [50–52]. Efficient protein degradation relies on proper positioning of autophagosomes and lysosomes[53]. Silencing of *NINL* results in impaired autophagic flux [54]. Impairment of molecular motors, stalled lysosomal motility, enhanced autophagic flux, and slowed cargo degradation have been described in *MAPT* mutant neurons [42, 55–58].

We propose that *NINL* overexpression reduces tau seeding by promoting protein clearance via the autophagy-lysosome pathway. HEK293T cells were transiently transfected with *NINL* WT to evaluate how *NINL* overexpression impacts components of the autophagy-lysosome pathway (**Figure 3A**). We used an acidotropic probe, LysoTracker Red, which accumulates in acidic vesicles, such as the lysosome, to evaluate the impact of *NINL* expression on lysosomes. Flow cytometry analyses revealed that LysoTracker intensity was significantly increased in cells overexpressing *NINL* compared to vector controls (**Figure 3B-C**). An increase in LysoTracker intensity could reflect a change in lysosomal load, acidity, or abundance. To determine whether increased *NINL* expression alters the acidity of lysosomal vesicles, we used a pH sensitive dye, LysoSensor Green (**Supplemental Figure 1A**). LysoSensor levels were similar in cells overexpressing *NINL* and vector controls (**Supplemental Figure 1B**).

**Figure 3.**
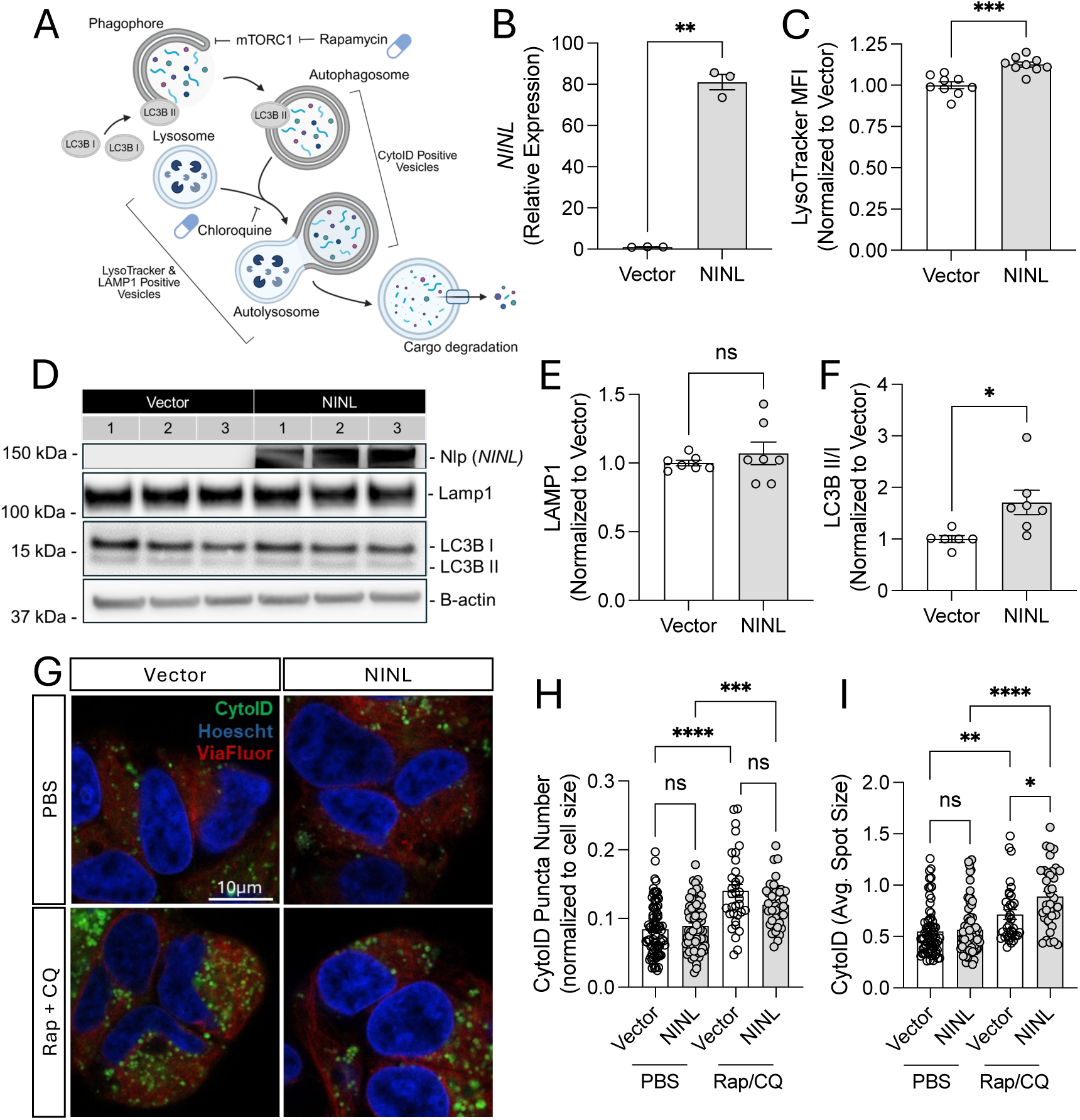
*NINL* expression enhances the autophagy-lysosome pathway. HEK293T cells were transiently transfected with pCMV6 NINL-myc plasmid or vector control plasmid for 48 hours. A. Schematic of the components of the autophagy-lysosome pathway that were evaluated. B. qPCR for *NINL* RNA. **, p = 0.0022. Student’s t-test. C. LysoTracker Median Fluorescence Intensity (MFI) quantified by flow cytometry plotted relative to vector control. ***, p = 0.0001. Student’s t-test. D. Representative immunoblot for Nlp (*NINL*), Lamp1, LC3B, and B-actin. E. Immunoblot quantification of Lamp1 protein levels normalized to vector controls. Ns, not significant. Student’s t-test. F. Immunoblot quantification of LC3B II/I ratio normalized to vector controls. *, p = 0.0224. Student’s t-test. G. Representative images from live imaging with CytoID. Both vector and *NINL* expressing cells were treated with rapamycin and chloroquine for 6 hours prior to imaging as a positive control. CytoID (autophagic vesicles) in green, Hoescht (nuclie) in blue, and ViaFluor (cytoskeleton) in red. Scale bar, 10μm. H. Quantification of the number of CytoID puncta per cell, normalized to the cell size (34-80 cells quantified per condition). ***, p = 0.0002; ****, p < 0.0001. Two-way ANOVA. I. Quantification of the average CytoID puncta size per cell. ****, p < 0.0001; **, p = 0.0048; *, p = 0.0151. Two-way ANOVA. Graphs represent mean ± SEM. Results represent 3 independent experiments with 2-3 replicates per condition in each experiment.

Thus, *NINL* overexpression does not cause hyperacidification of lysosomes. We next evaluated lysosomal abundance using immunoblotting for a lysosomal membrane marker, LAMP1 (**Figure 3A**). *NINL* overexpression resulted in similar levels of LAMP1 protein levels compared to vector controls (**Figure 3D-E**), suggesting neither lysosomal biogenesis nor mass is altered. Together, these findings suggest that *NINL* overexpression modulates lysosomal functional properties, without shifting basal lysosomal acidity or lysosomal abundance.

### 3.5 NINL alters autophagy

The observed increase in lysosomal functional properties, reflected by higher LysoTracker signal, without changes in lysosomal pH (LysoSensor) or overall lysosome abundance (LAMP1) could reflect altered cargo trafficking through the lysosomal system. Given that lysosomes represent the terminal compartment of the autophagy-lysosome pathway, an increase in lysosomal load could point to changes in autophagic input (**Figure 3A**). Thus, we next examined markers of autophagy to determine whether *NINL* expression alters autophagosomes. Initiation of autophagy involves the conversion of LC3BI to LC3BII, which integrates into the membrane of the forming phagophore and is ultimately degraded after autophagolysosomes undergo cargo degradation[59, 60]. Thus, ratios of LC3BII/I can provide insight into the initiation and completion of autophagy. The ratio of LC3B II/I was significantly elevated in HEK293T cells overexpressing *NINL* compared to vector control (**Figure 3D** and **3F**). Thus, at baseline, *NINL* likely modulates autophagy initiation or LC3 processing. We next sought to determine whether *NINL* expression alters autophagosome number or size using the cationic amphiphilic tracer, CytoID, that selectively labels autophagic vesicles. HEK293T cells overexpressing *NINL* or a control vector produced a similar number of autophagic vesicles (**Figure 3G**-**3H**) that were similar in size (**Figure 3G** and **3I**) under basal conditions. This finding, along with the change in LC3B II/I, suggests that *NINL* expression alters autophagy initiation or LC3 processing without expanding the autophagic vesicle pool under basal conditions [59]. To directly examine the impact of *NINL* on autophagosome dynamics, we used rapamycin to induce autophagy and chloroquine to block autophagosome-lysosome fusion in cells expressing control and *NINL* plasmids (**Figure 3G-I**). CytoID-positive vesicle number and size increased in vector and *NINL* expressing cells upon treatment with rapamycin and chloroquine compared to vehicle controls (**Figure 3G-I**). In rapamycin and chloroquine treated cells, *NINL* overexpression resulted in significantly larger CytoID-positive vesicles without a change in CytoID-positive vesicle number compared to vector control (**Figure 3G-I**). Taken together, these results suggest that increasing *NINL* regulates autophagosome maturation or expansion rather than vesicle biogenesis.

### 3.6 *NINL* promotes proteolytic activity

The observation that *NINL* increases lysosomal properties (higher LysoTracker) and enhances autophagosome maturation/expansion (increased LC3 lipidation and autophagic vesicle size with autophagy induction) points to activation of structural components of the degradative pathway without revealing the functional impact on degradative capacity. We next assessed lysosomal proteolytic activity to determine whether *NINL* promotes effective lysosomal degradation or leads to accumulation of undegraded material despite structural expansion. DQ-BSA is a fluorogenic protease substrate that is taken up by endocytosis and degraded by proteases, at which point it fluoresces [59]. Live imaging was performed in HEK293T cells overexpressing *NINL* or vector controls treated with DQ-BSA Red. Cells were imaged every two hours for 48 hours. We observed a similar increase of DQ-BSA signal in control and *NINL* expressing cells for the first 10 hours after treatment (**Figure 4A**). Beginning at 12 hours, *NINL* expressing cells produced significantly more DQ-BSA signal compared with cells expressing a vector control (**Figure 4A-B**). DQ-BSA signal is mediated by substrate uptake and proteolytic degradation. To further evaluate the impact of *NINL* expression on proteolytic activity, a membrane permeable fluorogenic substrate, Magic Red, was used[61]. Hourly live imaging revealed that HEK293T cells overexpressing *NINL* produced significantly more Magic Red signal than vector controls (**Figure 4C-D**). Thus, we show that *NINL* promotes proteolytic activity.

**Figure 4.**
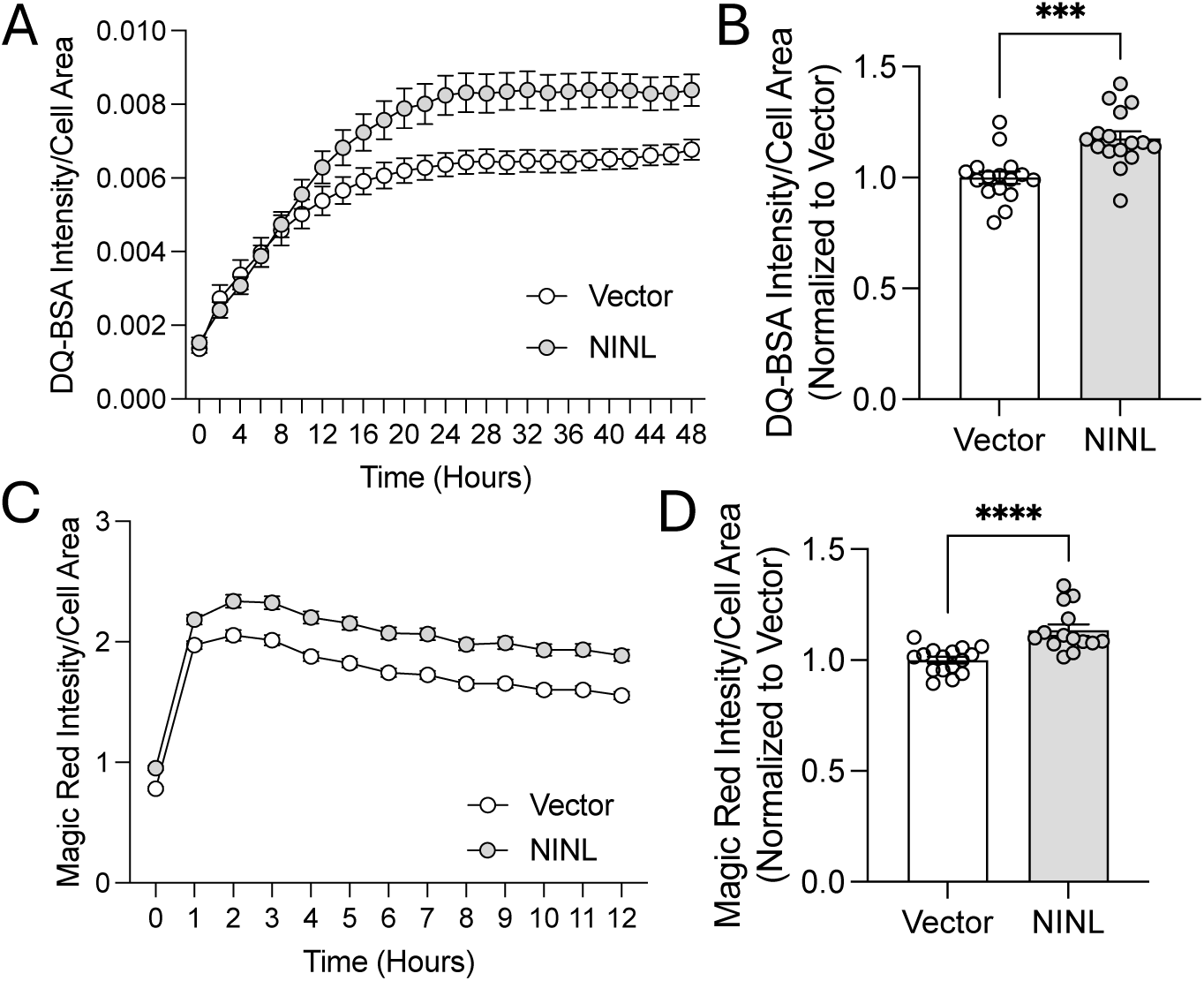
Increased *NINL* expression alters protease activity. HEK293T cells were transiently transfected with pCMV6 NINL-myc plasmid or vector control for 48 hours. A-B. Immediately after treatment with DQ-BSA Red, wells (96 well plate) were imaged every 2 hours at 10x (5 images per well). A. Representative graph of DQ-BSA integrated intensity over cell area across a 48-hour period. B. DQ-BSA integrated intensity over cell area normalized to vector controls at 48 hours. Each datapoint is representative of the average of 5 pictures per well. ***, p = 0.0002. Student’s t-test. C-D. Immediately after treatment with Magic Red, wells (96 well plate) were imaged every hour at 10x (5 images per well). C. Representative graph of Magic Red integrated intensity over cell area across a 12-hour period. D. Magic Red integrated intensity over cell area normalized to vector controls at 12 hours. Each datapoint is representative of the average of 5 pictures per well. ****, p < 0.0001. Student’s t-test. Graphs represent mean ± SEM. Results represent 2 independent experiments with 6-8 replicates per condition in each experiment.

### 3.7 *NINL* variants block protective effects on tau seeding

Our findings suggest that *NINL* overexpression reduces tau seeding via the autophagy-lysosome pathway by promoting autophagosome compartment abundance and degradative activity. Four *NINL* variants were identified in FTD cases and were absent from AD and control subjects (**Figure 5A; Supplemental Table 2**). *NINL* p.T275fs and p.R1202fs were predicted to be the most deleterious with high CADD scores (22.5 and 26.6, respectively), suggesting the variants are damaging. Both variants are predicted to cause a frameshift that results in an early stop codon and truncation of the Nlp protein. To determine whether *NINL* p.T275fs and p.R1202fs alter the Nlp protein, we mutagenized the *NINL* WT plasmid to introduce each variant (**Figure 5A**). Transient overexpression of plasmids containing *NINL* WT, p.T275fs, and p.R1202fs or vector control in HEK293T cells illustrated that the variants indeed reduced Nlp expression compared to *NINL* WT (**Figure 5B, Supplemental Figure 2**).

**Figure 5.**
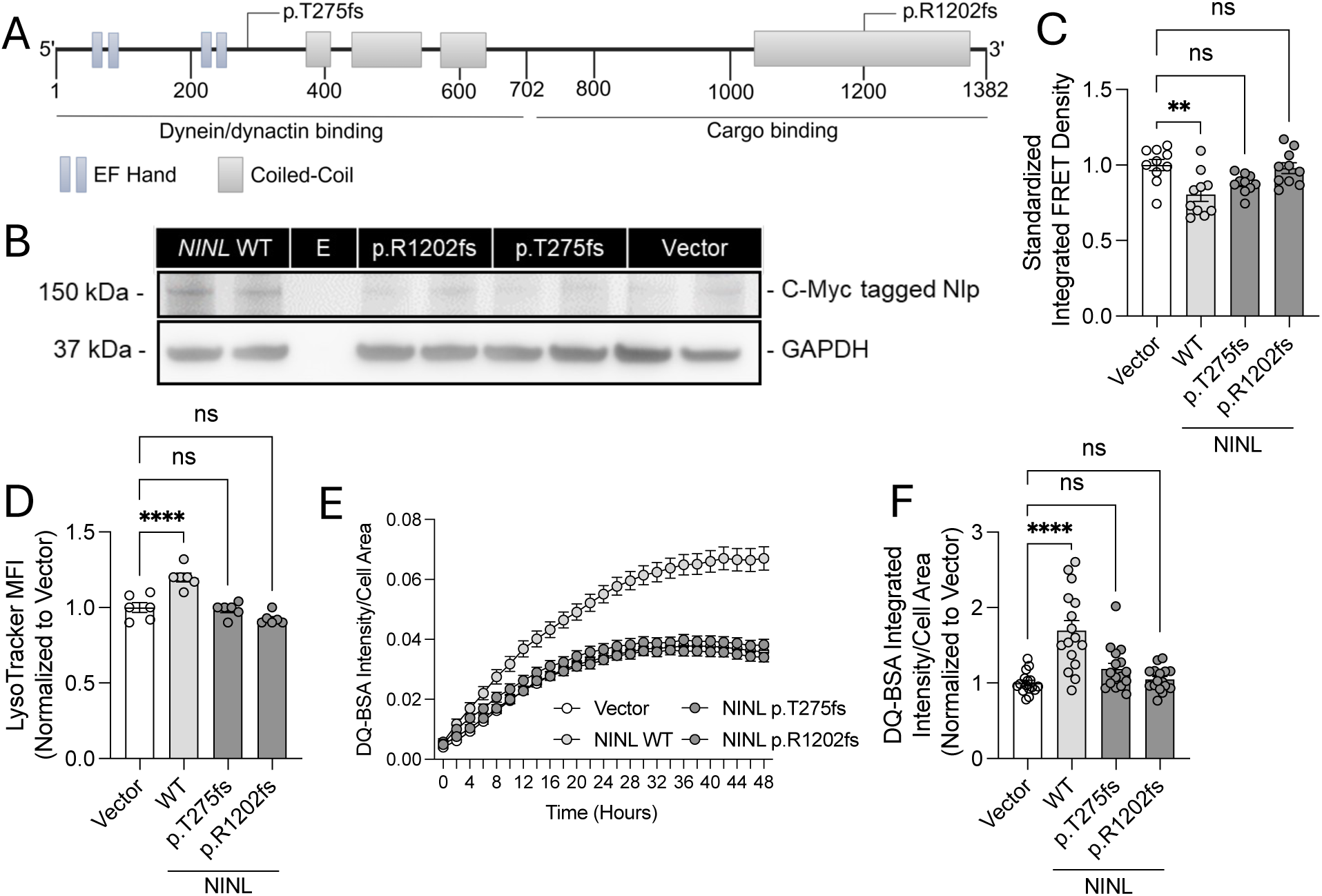
The protective effect of increased *NINL* expression on tau seeding is lost in *NINL* variants. Cells were transiently transfected with vector control, NINL WT, NINL p.T275fs, or NINL p.R1202fs. A. Schematic of the *NINL* gene and the variants identified in this study. B. Representative immunoblot for C-Myc tagged Nlp (*NINL*). C. Integrated FRET Density (FRET MFI x percent FRET positive cells) normalized to vector controls. **, p = 0.0014. One-way ANOVA with Dunnett correction. D. LysoTracker Median Fluorescence Intensity (MFI) quantified by flow cytometry plotted relative to vector control. ****, p < 0.0001. Student’s t-test. E. Representative graph of DQ-BSA integrated intensity over cell area across a 48-hour period. F. DQ-BSA integrated intensity over cell area normalized to vector controls at 48 hours. Each datapoint is representative of the average of 5 pictures per well. ****, p < 0.0001. Student’s t-test. Graphs represent mean ± SEM. Results represent 2 independent experiments with 5 replicates per condition in each experiment.

We next sought to determine whether *NINL* variants alter tau seeding. Consistent with our prior experiments, expression of *NINL* WT in HEK293T tau biosensor cells resulted in a significant reduction of tau seeding (**Figure 5C**). However, expression of *NINL* p.T275fs and p.R1202fs in tau biosensor cells failed to reduce tau seeding (**Figure 5C**). Thus, FTD variants in *NINL* eliminate the protective effect of *NINL* on tau seeding.

*NINL* variants may fail to reduce tau seeding by disrupting the effects of *NINL* in the autophagy-lysosome system. Consistent with our prior experiments, expression of *NINL* WT in HEK293T cells resulted in a significant increase in LysoTracker intensity, pointing to increased lysosomal functional properties (**Figure 5D**). However, expression of *NINL* p.T275fs and p.R1202fs failed to alter LysoTracker intensity compared to vector controls (**Figure 5D**). To understand the impact of *NINL* p.T275fs and p.R1202fs on protease activity, HEK293T cells were transiently transfected with *NINL* WT, p.T275fs, p.R1202fs, or vector control, treated with DQ-Red BSA, and imaged every two hours for 48 hours (**Figure 5E**). Cells expressing *NINL* p.T275fs and p.R1202fs behaved similarly to vector controls, exhibiting less protease activity than *NINL* WT (**Figure 5E-F**). Together, we discovered that *NINL* p.T275fs and p.R1202fs block the protective effect of *NINL* WT on tau seeding by failing to promote lysosomal protease activity.

## 4. Discussion

Proteostasis dysfunction is a hallmark of FTD and AD, yet the genetic factors and molecular pathways that disrupt protein homeostasis remain poorly understood. Here, we integrated human genetics, transcriptomics, and mechanistic studies to define proteostasis network components contributing to tauopathy. Using genome sequencing from unrelated FTD, early-onset AD, and control subjects, we identified 18 proteostasis network genes harboring 75 rare, predicted-damaging variants enriched in FTD and/or AD. These genes span diverse proteostasis functions, including autophagy-lysosome regulation, ubiquitin-proteasome activity, and chaperone regulation. Many of these proteostasis network genes were differentially expressed in *MAPT* mutant neurons and dysregulated in FTLD-tau, AD, and PSP brains, demonstrating convergence of genetic and transcriptional perturbations. A single gene, *NINL*, emerged as consistently upregulated across *MAPT* mutant neurons, FTLD-tau, AD, and PSP brains. Functional studies demonstrated that *NINL* overexpression reduces tau seeding, increases LC3B lipidation, enlarges autophagic vesicles during flux blockade, elevates LysoTracker signal without altering lysosomal pH or mass, and enhances proteolytic activity, supporting a role for *NINL* in promoting autophagosome maturation and lysosomal degradation. FTD-enriched *NINL* variants (p.T275fs, p.R1202fs) disrupted Nlp protein expression and abolished *NINL*-mediated reductions in tau seeding and lysosomal protease activity. Together, these findings identify a set of proteostasis genes genetically and transcriptionally linked to neurodegeneration and reveal *NINL* as a novel regulator of tau aggregation, potentially upregulated as an adaptive response to proteotoxic stress.

Proteostasis, the coordinated regulation of protein synthesis, folding, trafficking, and degradation, is essential for neuronal health, and its breakdown is a defining feature of neurodegenerative diseases such as FTLD-tau and AD. Since these disorders typically arise in mid-to-late adulthood, proteostasis decline has often been viewed as an age-related vulnerability that acts as a secondary hit in cells already burdened by misfolded proteins. However, emerging evidence indicates that defects in core proteostasis machinery can occur much earlier, prior to clinical onset, and may actively contribute to the initiation or acceleration of tau pathology [8–11, 14–16, 19, 20].

Our sequencing analyses identified rare variants enriched in FTD and AD across multiple functional classes of proteostasis genes, including those encoding lysosomal enzymes, trafficking regulators, and components of the autophagic machinery. This suggests that genetic variation in protein clearance pathway genes is relevant to neurodegeneration and, specifically, that rare variation in these genes may directly affect degradation capacity by altering distinct but converging proteostasis nodes. Transcriptomic analyses further demonstrated that these proteostasis genes are consistently dysregulated in iPSC-derived neurons carrying *MAPT* mutations, which model early disease processes, and in FTLD-tau, AD, and PSP brains, which represent end-stage pathology. This convergence across early, pre-aggregation neuronal states and advanced human brain pathology suggests that proteostasis genes harboring rare variants represent components of degradation pathways systematically stressed across disease stages. Directionality of the proteostasis gene expression changes (upregulation vs. downregulation) may reflect compensatory responses, pathway overload, or direct mutation effects. Additional functional analyses are required to elucidate these mechanisms. Together, this convergence across genetics, patient-derived neurons, and diseased brains supports a model in which both inherited variation and disease-driven cellular stress perturb common proteostasis pathways, underscoring the biological relevance of these genes to neurodegenerative processes.

Among the candidate genes, *NINL* (*Ninein-like*) emerged as a compelling regulator of proteostasis given its consistent upregulation across *MAPT* mutant neurons and tauopathy brains. *NINL* encodes the motor associated protein Nlp which is canonically known for its role in the centrosome[47–49]. In neurons, Nlp supports vesicle trafficking by interacting with motor complexes, such as dynein and dynactin [50–52], placing it at a strategic node for coordinating autophagic and lysosomal transport [54].

Functional studies revealed that *NINL* overexpression enhances autophagic degradation capacity: it increases LC3BII/I ratio at baseline, enlarges CytoID-positive vesicles under rapamycin and chloroquine–induced flux blockade, and elevates LysoTracker signal without altering lysosomal abundance or pH. *NINL* also enhanced lysosomal functional properties and proteolytic activity, reflected by increased LysoTracker retention and accelerated degradation of DQ-BSA and Magic Red substrates. These findings demonstrate that *NINL* strengthens the degradative arm of the autophagy-lysosome pathway. Together, these data support a model in which *NINL* promotes effective clearance of aggregation-prone proteins by improving autophagy–lysosome coupling and degradative efficiency.

Importantly, we show that *NINL* reduces tau aggregation. Two FTD-enriched *NINL* frameshift variants abolished the protective effects of *NINL* on tau seeding and failed to enhance lysosomal proteolysis. These variants reduced Nlp protein expression, indicating loss-of-function effects. Their inability to reduce tau seeding or promote proteolytic activity suggests that the genetic disruptions in *NINL* observed in FTD cases may impair the cell’s capacity to counteract pathological tau accumulation. This provides direct functional evidence linking *NINL* genetic variation to altered proteostasis, offering mechanistic insights into how rare variants might contribute to disease. The combined genetic, transcriptomic, and functional evidence supports a model in which *NINL* acts as a molecular amplifier of degradative capacity. *NINL* upregulation observed in human disease tissue may therefore represent an adaptive, compensatory response to rising proteotoxic burden, aimed at maintaining proteostasis in the face of accumulating tau pathology. Similar compensatory upregulation has been documented for other autophagy and lysosomal genes in early or presymptomatic disease[62], highlighting the dynamic nature of proteostasis responses.

Together, this work broadens the spectrum of proteostasis components implicated in disease, demonstrating that rare variants and disease-driven dysregulation extend beyond canonical autophagy and lysosome genes. Our findings reveal a coordinated pattern of genetic burden, dysregulation of gene expression, and impaired proteostasis involving a subset of proteostasis genes in FTD and AD. By identifying *NINL* as a regulator of tau aggregation and lysosomal degradation, we highlight a previously uncharacterized molecular contributor to disease pathophysiology. These results suggest that enhancing *NINL* function or strengthening downstream trafficking and degradative pathways may represent promising therapeutic strategies for tauopathies.

Several limitations warrant consideration. While rare variant enrichment highlights genes of potential importance, further functional validation of these variants will be required to establish their disease risk effect. Transcriptomic datasets lack single-cell resolution, limiting our ability to attribute expression changes to specific cell types or disease stages. Finally, while iPSC-derived neurons and overexpression assays capture many critical aspects of human proteostasis networks, *in vivo* studies will be needed to determine whether *NINL* modulation alters tau propagation, neuronal survival, and cognitive outcomes.

In summary, our integrative approach demonstrates that rare genetic variation, transcriptional dysregulation, and functional perturbation of proteostasis genes converge on pathways critical for tau aggregation. We identify NINL as a mechanistically relevant regulator of autophagic degradation and tau aggregation and propose that its upregulation in disease reflects an adaptive attempt to bolster proteostasis capacity. These insights advance our understanding of tau clearance mechanisms and nominate proteostasis pathways as actionable targets for therapeutic intervention in FTD and AD.

## Acknowledgements

This work was supported by access to equipment made possible by the Hope Center for Neurological Disorders, the Department of Pathology and Immunology, Center of Cellular Imaging, and the Department of Psychiatry at Washington University School of Medicine. Live imaging experiments were performed using the Washington University Center for Cellular Imaging (WUCCI) supported by Washington University School of Medicine, The Children’s Discovery Institute of Washington University and St. Louis Children’s Hospital (CDI-CORE-2015-505 and CDI-CORE-2019-813) and the Foundation for Barnes-Jewish Hospital (3770 and 4642). Confocal data was generated on a Nikon AX-R Confocal Microscope which was purchased with support from the Office of Research Infrastructure Programs (ORIP), a part of the NIH Office of the Director under grant OD030233. FRET flow cytometry experiments were performed using the Washington University Department of Pathology and Immunology Flow Cytometry Core. Diagrams were generated using BioRender.com. The content of this publication is solely the responsibility of the authors and does not necessarily represent the official views of the NIH.

## Disclosures

CMK serves as an advisor for Eisai Co. Ltd and Synapticure Inc. JSY serves on the scientific advisory board for the Epstein Family Alzheimer’s Research Collaboration and the Charleston Conference on Alzheimer’s Disease and is the editor-in-chief of *npj Dementia*.

## Sources of Funding

Funding provided by the National Institutes of Health (NS123985 [CMK, JSY], AG066444 [CMK], NS110890 [CMK], UL1TR002345, T32AG058518 [SKS], AG062588 [JSY], AG057234 [JSY], AG062422 [JSY], AG019724 [JSY], AG079774 [JSY]), Rainwater Charitable Foundation (CMK, JSY), Hope Center for Neurological Disorders (CMK), Global Brain Health Institute (JSY), and the Mary Oakley Foundation (JSY).

## Consent Statement

All participants provided informed consent for their data to be used in this study. The study was approved by the institutional review board of Washington University School of Medicine in St. Louis.

## Authors’ Contributions

Designed experiments: SKS, JSY, CMK. Performed and analyzed experiments: SKS, GH, JNC, MAM, PAC, KM, ES, GG, JAM, AK, JSY, CMK. Provided funding: AK, JSK, CMK. Wrote the manuscript: SKS, CMK. Revised and approved manuscript: SKS, GH, JNC, MAM, AC, KM, ES, GG, JAM, AK, JSY, CMK.

## Notes

### Competing Interest Statement

The authors have declared no competing interest.

