## Supplemental Figure 1 for "Integrative Genomic and Functional Analyses Reveal NINL as a Modulator of Tau Aggregation"

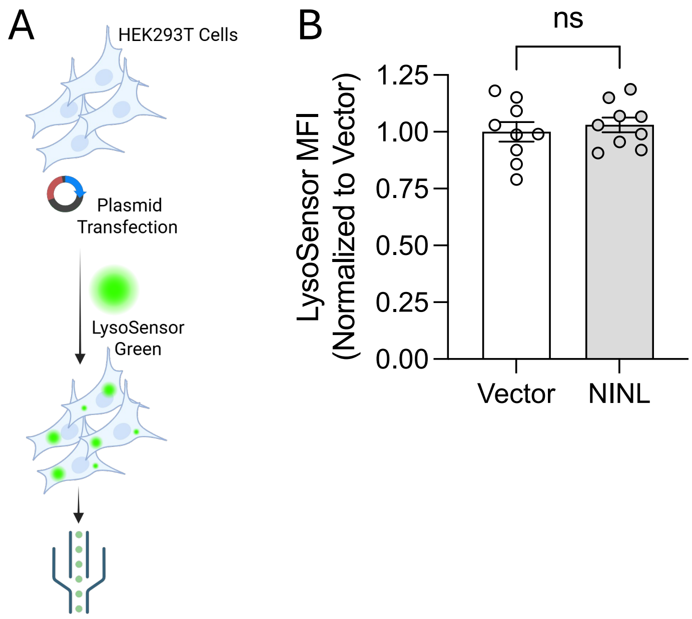


**Supplemental Figure 1. *NINL* expression does not alter lysosomal acidity.** A. Schematic of LysoSensor flow experiment in HEK293T cells. B. LysoSensor MFI (Median Fluorescence Intensity) quantified by flow cytometry plotted relative to vector control. Ns = non-significant. Student's t-test. Graphs represent mean ± SEM. Results represent 3 independent experiments with 2-3 replicates per condition in each experiment.
