## Supplemental Figure 2 for "Integrative Genomic and Functional Analyses Reveal NINL as a Modulator of Tau Aggregation"

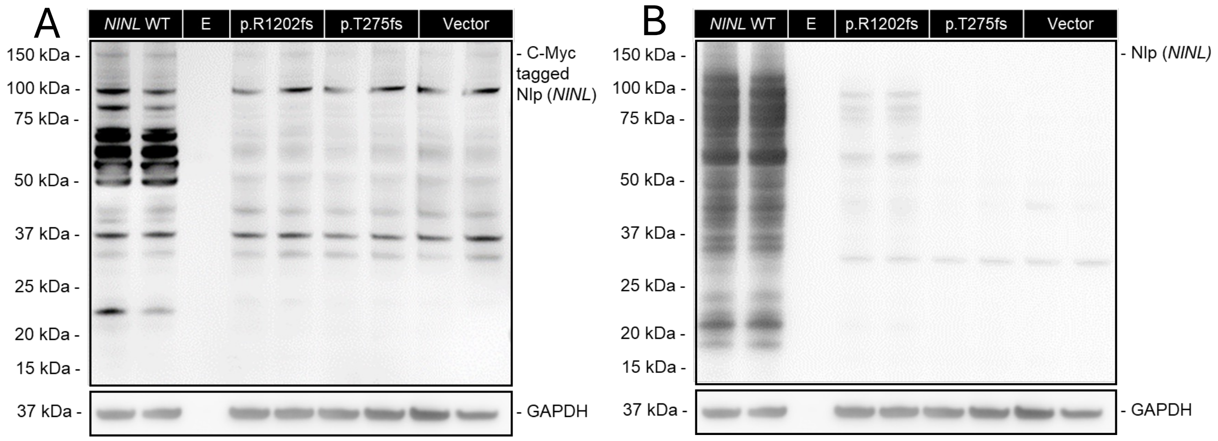


**Supplemental Figure 2. Rare, frame shift *NINL* variants disrupt Nlp protein.** A. Representative immunoblot for 9E10, which detects the C-terminal c-myc tag on Nlp. B. Representative immunoblot for Nlp. Results representative of 1 experiment with 2 replicates per condition.
